# Integrated Human Transcriptomics Identifies Fallopian Tube Progenitors as Plausible Precursors of High-Grade Serous Ovarian Cancer

**DOI:** 10.1101/2025.09.26.678859

**Authors:** Qian Li, Keren Cheng, Lili Sun, Wei Yan

## Abstract

High-grade serous ovarian cancer (HGSOC) is the most lethal gynecologic malignancy, yet its earliest cellular precursors remain incompletely defined. Here, we integrate bulk and single-nucleus RNA sequencing of human fallopian tubes, ovaries, and primary HGSOC tumors to resolve the epithelial hierarchy of the fallopian tube and examine its relationship to malignant states. We identify a rare LGR5⁺/PGR⁺ basal population as putative epithelial stem cells and delineate a large pool of OVGP1⁺/RNPC3⁺ progenitors that give rise to both secretory and ciliated lineages. By jointly analyzing normal and malignant transcriptomes, we find that these progenitor populations show the strongest transcriptional and developmental continuity with multiple HGSOC cell states, suggesting that they represent the epithelial compartments most plausibly poised for oncogenic divergence. Transcription factor network reconstruction highlights SPDEF as a key regulator of progenitor identity and NR2F6 as a candidate oncogenic driver, while inferred copy number variations and microenvironmental collagen-integrin signaling further illuminate the genomic and stromal factors that consolidate malignant fate. Together, these findings refine the cellular landscape of the human fallopian tube and identify progenitor-enriched molecular features that could inform earlier detection, improved risk stratification, and new preventive strategies for women at risk of developing HGSOC.

**One sentence summary:** Fallopian tube progenitor cells represent a key intermediate state linking normal epithelial hierarchy to the earliest stages of high-grade serous ovarian cancer development.

## INTRODUCTION

High-grade serous ovarian carcinoma (HGSOC) is the most common and lethal subtype of ovarian cancer, accounting for more than 70% of ovarian cancer deaths and associated with a 10-year survival rate below 30% (*1, 2*). For decades, HGSOC was thought to arise from the ovarian surface epithelium (OSE), but extensive molecular and pathological evidence has fundamentally reshaped this view over the past two decades (*3, 4*). HGSOC more closely resembles Müllerian duct-derived epithelia of the fallopian tube than the mesothelial-like OSE, based on its transcriptional, proteomic, and morphological features (*5, 6*). These observations have led to the now widely supported hypothesis that the fallopian tube epithelium (FTE), particularly the distal fimbrial region, represents the most plausible origin for many HGSOCs (*7, 8*).

Compelling support for this hypothesis comes from serous tubal intraepithelial carcinomas (STICs), early precursor lesions that harbor TP53 mutations identical to those in matched invasive tumors (*9, 10*). STICs are frequently found in the fimbriae of high-risk BRCA1/2 mutation carriers and in patients with sporadic HGSOC (*11–14*). While these lesions strongly link the distal fallopian tube to HGSOC pathogenesis, they do not identify which epithelial populations within the tube are most susceptible to transformation, nor how these populations depart from normal differentiation trajectories. Historically, although mature secretory cells have been proposed as likely precursors, recent single-cell studies reveal greater epithelial complexity (*15–18*), suggesting that ciliated-lineage or transitional states may also contribute to malignant evolution.

The fallopian tube consists of four anatomical segments: the infundibulum, ampulla, isthmus, and the uterotubal junction, each exposed to distinct physiological environments. The distal infundibulum and fimbria, positioned adjacent to the ovary, experience repeated bursts of inflammatory and oxidative stress during ovulation, necessitating frequent epithelial repair (*19–22*). Early histological studies described small, round “basal” or “intercalary” cells as potential stem cells capable of generating both ciliated and secretory lineages (*20, 23*), yet their molecular identity remains uncertain. Advances in single-nucleus RNA sequencing (snRNA-seq) offer a way to clarify such lineage relationships in human tissues, as this approach avoids dissociation-induced artifacts, accommodates frozen clinical samples, and produces high-resolution transcriptomic data suiData File for integration with pathology (*24–27*).

Understanding the epithelial hierarchy of the human fallopian tube and identifying the populations capable of initiating HGSOC have remained challenging due to limited access to healthy human tissues, a lack of definitive stem/progenitor markers, and the absence of lineage-tracing models in women. High-resolution transcriptomic technologies now allow for reconstruction of epithelial differentiation trajectories and direct comparison of normal epithelia with malignant states in primary tumors. In this study, we integrate bulk RNA sequencing of normal fallopian tubes, ovaries, and HGSOC samples with snRNA-seq of cancer-free fallopian tubes and primary HGSOC tissues. We define a stem-progenitor hierarchy within the FTE, identifying rare LGR5⁺/PGR⁺ basal cells as putative stem cells that give rise to a large and widely distributed pool of OVGP1⁺/RNPC3⁺ progenitor cells. By jointly analyzing normal and malignant transcriptomes, we show that these progenitors exhibit the strongest transcriptional and developmental continuity with multiple malignant trajectories, making them the epithelial populations most poised for oncogenic deviation. We further delineate transcription factor networks, copy-number alterations, and microenvironmental signaling pathways that accompany progression from normal epithelial differentiation to malignant transformation. Rather than asserting a definitive cell of origin, our findings provide a mechanistic and spatial framework indicating that fallopian tube progenitors are the most biologically plausible intermediates capable of evolving into HGSOC under oncogenic, inflammatory, or microenvironmental stress. These insights refine our understanding of epithelial organization in the fallopian tube and illuminate the earliest cellular and molecular events that may underlie HGSOC development, with implications for early detection, risk assessment, and targeted prevention.

## RESULTS

### Dual origins of HGSOC identified by bulk RNA-seq

High-grade serous ovarian cancer (HGSOC) has been proposed to arise from either the fallopian tube epithelium or the ovarian surface epithelium (*11, 28, 29*). To evaluate these possibilities, we performed bulk RNA-seq to examine transcriptomic relationships among normal fallopian tubes, normal ovaries, and HGSOC tumors (Fig. 1A). Twenty-six human specimens were analyzed, including four cancer-free ovaries, four cancer-free fallopian tubes dissected into three anatomical segments (fimbriae/infundibulum, ampulla, and isthmus), and ten HGSOC tumors. Cancer-free tissues were collected from women undergoing surgery for benign uterine conditions, whereas tumor samples were obtained from patients pathologically diagnosed with epithelial ovarian cancer and subsequently confirmed as HGSOC (Table S1).

**Figure 1.**
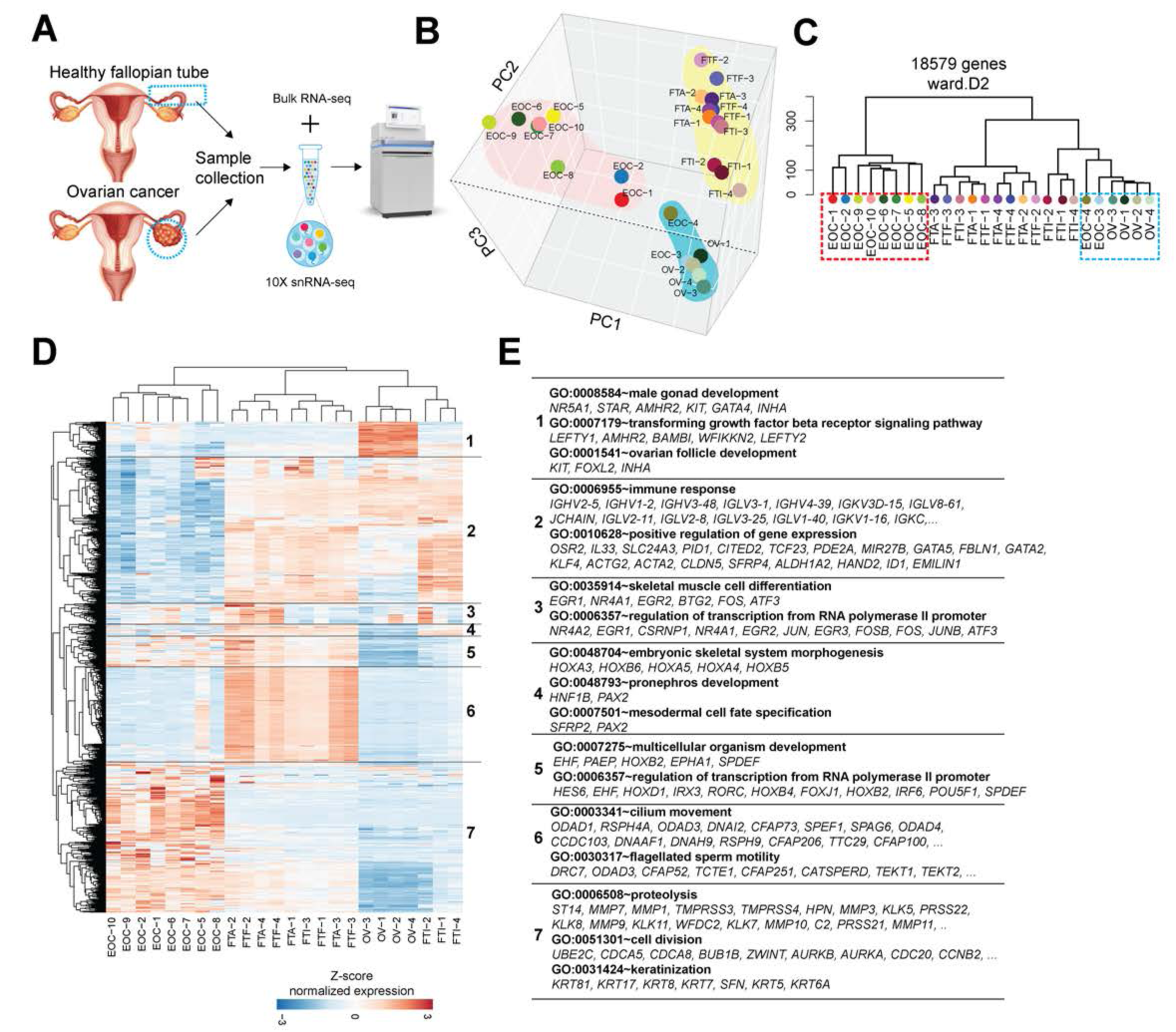
Transcriptomic relationships among human fallopian tubes, ovaries, and high-grade serous ovarian cancer. **(A)** Overview of human samples used for bulk RNA-seq, including cancer-free ovaries, cancer-free fallopian tube segments (isthmus, ampulla, fimbria), and high-grade serous ovarian cancer (HGSOC) tumors. **(B)** Principal component analysis of bulk RNA-seq data showing three distinct clusters corresponding to ovary, fallopian tube, and HGSOC. **(C)** Hierarchical clustering and heatmap of pairwise Pearson correlations identifying two HGSOC subgroups: fallopian tube–like (EOC_F) and ovary-like (EOC_O). **(D)** Heatmap of seven major gene sets differentially enriched among ovary, fallopian tube, and EOC_F samples. Representative genes for each set are shown. See also **Data File S1**.

Sequencing produced 38-65 million high-quality reads per sample (Fig. S1A). Principal component analysis revealed three clearly separated clusters corresponding to ovary, fallopian tube (FT), and HGSOC tissues (Fig. 1B). Pearson correlation analysis further highlighted the transcriptional distances among these groups. Notably, eight HGSOC samples clustered with fallopian tube tissues, whereas two samples (EOC-3 and EOC-4) clustered with ovaries, suggesting distinct tissue affinities. Hierarchical clustering confirmed this separation, allowing classification of EOC-3 and EOC-4 as ovary-like HGSOC (EOC-O) and the remaining eight tumors as fallopian tube-derived HGSOC (EOC-F) (Fig. 1C; Fig. S1B). Differential gene expression analysis revealed that fallopian tube-enriched genes, including *FOXJ1, SOX17, PODXL, MUC16*, and *WFDC2,* were markedly elevated in EOC-F tumors. In contrast, ovarian marker genes, including *FOXL2, ARX, KIT, STAR*, and *WFIKKN2,* were predominantly expressed in EOC-O tumors (Fig. S1C). Gene Ontology (GO) analysis showed enrichment of cell division, cell adhesion, and spermatogenesis pathways in EOC-F, whereas EOC-O exhibited signatures associated with gonadal development and WNT signaling (Fig. S1C). Together, these results support a dual-origin model of HGSOC, with the majority of tumors in our cohort derived from the fallopian tube.

Clustering of all differentially expressed genes across the ovary, FT, and EOC-F samples identified seven major gene sets representing core transcriptional programs (Fig. 1D; Data File S1). Ovarian tissues showed enrichment for genes involved in TGF-β signaling and gonadal development (Geneset 1), whereas normal FT and ovary shared immune regulatory and transcriptional programs (Geneset 2). Additional gene sets reflected smooth muscle components (Geneset 3), FT-specific developmental signatures, including *PAX2* and *HOX* family genes (Geneset 4), and genes upregulated in both FT and EOC-F, such as *SPDEF, POU5F1, FOXJ1, HOXB2*, and *HOXB4* (Geneset 5). Cilium assembly genes (Geneset 6) and cancer-associated pathways involving proliferation, metabolism, and keratinization (Geneset 7) were also distinctly represented. Pairwise comparisons among EOC-F, FT, and ovary samples (Fig. S1D–F) highlighted pathways involved in cell division, chemokine signaling (e.g., *CXCL9/10/11-CXCR3, CXCL6-CXCR2*), inflammatory responses (*IL1-IL1RAP, GAL-3, NR1H4*), apoptosis (*IL24, TNFAIP8*), and antigen presentation via MHC class II. Immune-related genes, such as *HLA-DMA* and *HLA-DRA*, which are key to T-cell recruitment and differentiation, were strongly expressed, consistent with the inflammatory and immune-rich microenvironment of HGSOC. These findings underscore the immunologic complexity of fallopian tube-derived tumors. Collectively, our bulk RNA-seq analyses establish that HGSOC comprises at least two transcriptionally distinct subtypes, one resembling fallopian tube epithelium and the other resembling ovarian tissue. The predominance of the FT-like subtype in our cohort further reinforces the fallopian tube as the principal tissue of origin for most HGSOCs.

### Identification of a large pool of progenitors capable of differentiating into secretory or ciliated cells in the fallopian tube epithelium

Although several recent single-cell studies have profiled human fallopian tube epithelium (FTE) (*21, 30–33*), the molecular identity of true FTE stem cells, the hierarchy of secretory cell subtypes, and the developmental relationship between mature secretory and ciliated cells remain incompletely defined. To address these uncertainties, we collected cancer-free fallopian tubes from three perimenopausal women undergoing hysterectomy with bilateral salpingo-oophorectomy for benign uterine conditions (Table S2). Each tube was dissected into fimbriae/infundibulum, ampulla, and isthmus, and single-nucleus RNA sequencing (snRNA-seq) was performed on nine samples.

Using the 10x Genomics platform, we obtained high-quality transcriptomes from 87,920 nuclei after initial quality control, with 36,000-63,000 mean reads per nucleus and approximately 2,000 genes detected per cell (Fig. S2A). After regressing out cell cycle effects, mitochondrial gene content, and library size, no cell cycle-driven clustering was observed (Fig. S2B), and all segments were evenly represented on UMAP, indicating minimal batch-related segregation (Fig. S2C). These results confirmed the robustness of the dataset. Initial clustering of all fallopian tube samples identified five major cell types: secretory cells, ciliated cells, smooth muscle cells, endothelial cells, and macrophages (Fig. S2D). Marker gene (Fig. S2E) and GO enrichment (Fig. S2F; Data File S2) analyses suggested that the secretory cell cluster contained the putative stem cell population. This cluster showed higher expression of genes associated with adult stem cell maintenance and epithelial progenitor states, including *SOX17, LGR5, TFCP2L1,* and *HES1*. Endothelial, smooth muscle, ciliated, and macrophage clusters each expressed expected lineage-specific signatures (Fig. S2F). The distribution of major epithelial populations across segments showed an increasing proportion of ciliated cells from the isthmus to the fimbriae, and the inverse pattern for secretory cells (Fig. S2G). To resolve epithelial heterogeneity at higher resolution, we reclustered 34,267 epithelial cells, yielding 12 distinct epithelial subclusters (Fig. 2A). Hierarchical clustering grouped these into five broader categories (Fig. 2B). Based on marker expression, cluster 12 was identified as the FTE stem cell population, defined by high expression of *LGR5, PGR, PODXL,* and *SOX17*. Clusters 2 and 6 expressed *PAX8, ZBTB16,* and *EGR1* and were annotated as mature secretory cells. Clusters 1, 4, 5, and 8 represented ciliated cells expressing *CAPS, CFAPs*, and dynein-associated genes. The remaining clusters, 3, 7, 9, 10, and 11, expressed *OVGP1, CRISP3, RNPC3,* and *BDP1* but lacked stem cell and mature cell markers, indicating they are likely progenitor cells (Fig. 2C).

**Figure 2.**
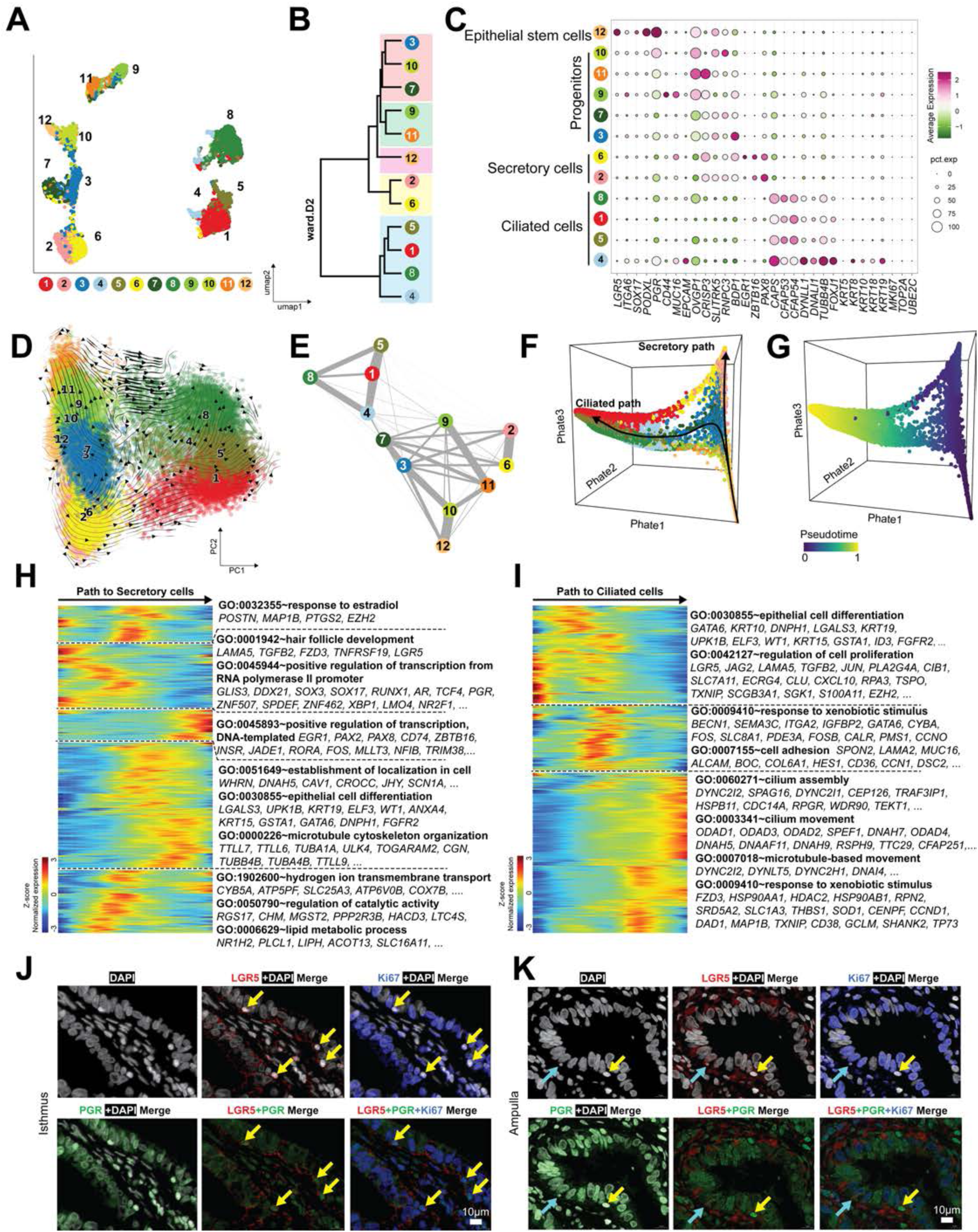
Cellular composition and differentiation hierarchy of the human fallopian tube epithelium revealed by snRNA-seq. **(A)** UMAP visualization of 34,267 epithelial nuclei from human fallopian tubes, identifying 12 epithelial subclusters. **(B)** Hierarchical clustering of subclusters grouped into five major epithelial categories. **(C)** Dot plot showing marker genes defining stem cells, progenitors, secretory cells, and ciliated cells. **(D)** RNA velocity fields projected onto UMAP indicate directional flow from stem cells toward either secretory or ciliated lineages. **(E)** PAGA graph depicting connectivity among epithelial states, placing stem cells at the root. **(F–G)** PHATE embeddings and pseudotime trajectories showing two major differentiation paths: secretory and ciliated. **(H–I)** Heatmaps of dynamic gene expression along secretory (H) and ciliated (I) trajectories. **(J–K)** Immunofluorescence localization of LGR5 and PGR in basal epithelial stem cells of the fallopian tube. Scale bars: 50 µm. See also **Figure S2** and **Data File S2–S4**.

RNA velocity analysis demonstrated that progenitor cells occupy transitional positions between stem cells and both mature lineages, with directional flow toward either secretory or ciliated fates (Fig. 2D). PAGA confirmed strong connectivity: stem cells (Cluster 12) positioned at the root, progenitors forming intermediate nodes, and mature secretory and ciliated cells at opposite termini (Fig. 2E). PHATE visualization revealed two branching developmental trajectories, supported by pseudotime inference (Fig. 2F–G). Marker genes such as *LGR5, PAX8, FOXJ1,* and *CFAP54* aligned with expected pseudotime patterns (Fig. S2H). Transcriptomic dynamics along the secretory trajectory showed gradual downregulation of stem cell markers (*LGR5, SPDEF, NR2F1, AR*) and transient induction of estradiol-responsive genes (*POSTN, MAP1B, PTGS2, EZH2*), followed by increased expression of mature secretory genes, including *EGR1, PAX2, PAX8, CD74,* and *ZBTB16* (Fig. 2H; Data File S3). Along the ciliated trajectory, early decreases in *LGR5* and *LAMA5* were followed by transient induction of xenobiotic response and adhesion-related genes, and terminal enrichment of CFAPs and dyneins associated with ciliogenesis (Fig. 2I; Data File S4). Quantification of epithelial subtypes revealed that progenitors comprise the largest epithelial population, ∼38% of all epithelial cells. Stem cells were concentrated in the isthmus and decreased toward the fimbriae, whereas progenitors remained consistently abundant across segments (Fig. S2I–J). Immunostaining supported these observations: TUBB4B⁺ ciliated cells increased from isthmus to fimbriae, while PAX8⁺ secretory cells decreased (Fig. S3A–B), matching snRNA-seq distributions. LGR5 and PGR co-localized in small round basal cells, confirming their identity as FTE stem cells (Fig. 2J–K; Fig. S3C). Stem cells were enriched in the isthmus and were rare in the fimbriae, whereas progenitors were abundant in all segments. Together, these data define a hierarchical epithelial organization in which rare LGR5⁺/PGR⁺ stem cells generate a large, widely distributed pool of OVGP1⁺ progenitors that can differentiate into either mature secretory or ciliated cells. This progenitor population likely represents the key transitional state underlying both normal epithelial turnover and vulnerability to malignant transformation.

### High inter-patient variability and heterogeneity in HGSOC cell composition

High-grade serous ovarian cancer (HGSOC) is characterized by substantial cellular complexity and inter-patient variability. To define the composition of malignant and non-malignant cell populations within HGSOC, we analyzed 15,322 nuclei from snRNA-seq datasets generated from four primary tumors (Fig. S3D). After removing batch effects and regressing out cell cycle and library size using LIGER (*34*), datasets showed no residual technical clustering (Fig. S3E), yet pronounced biological heterogeneity across patients remained evident (Fig. S3F–G). Unsupervised Louvain clustering resolved 13 transcriptionally distinct clusters (Fig. 3A). Hierarchical clustering grouped these into ten malignant cell clusters and three non-malignant clusters corresponding to macrophages, cancer-associated fibroblasts (CAFs), and endothelial cells (Fig. 3B). Malignant clusters represented five major cancer cell states: *proliferative, mesenchymal-like, ciliated-like, epithelial-to-mesenchymal transition (EMT), and MHC (immunoreactive)* (Fig. 3C). Cluster-specific marker analysis (Fig. 3D; Data File S5) showed that proliferative cancer cells strongly expressed genes regulating cell cycle progression and mitosis, consistent with a tumor stem-like or progenitor-like phenotype. Mesenchymal-like cancer cells expressed transcription factors linked to cellular plasticity and migration (*SOX17, MECOM, PAX8*) along with markers of matrix remodeling, suggesting a role in invasion and stromal engagement. Ciliated-like cancer cells expressed cilium assembly and motility genes, indicating partial retention of their fallopian tube lineage identity. EMT-like cancer cells upregulated genes associated with cell motility and extracellular matrix interactions, reflecting active epithelial-mesenchymal transition. MHC (immunoreactive) cancer cells displayed robust expression of antigen processing and MHC class II-related genes, underscoring their interaction with the immune microenvironment.

**Figure 3.**
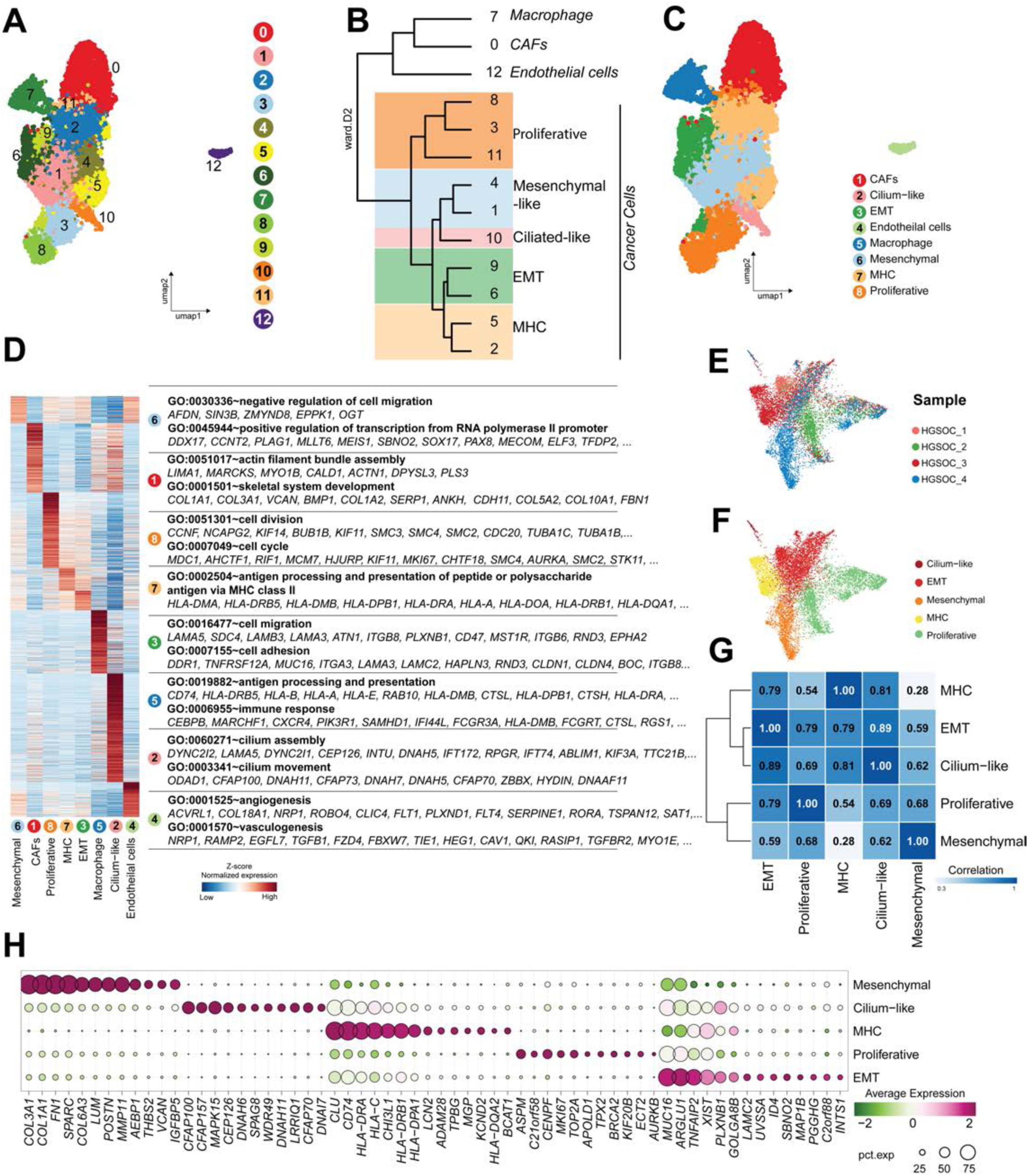
Cellular diversity and inter-patient heterogeneity in human HGSOC. **(A)** UMAP plot of 15,322 nuclei from four HGSOC tumors identifying 13 cellular clusters. **(B)** Dendrogram showing three non-malignant clusters (macrophages, cancer-associated fibroblasts (CAFs), endothelial cells) and five major HGSOC cell states. **(C)** Annotation of cancer subtypes: proliferative, mesenchymal-like, epithelial-to-mesenchymal transition (EMT), ciliated-like, and immunoreactive (MHC). **(D)** Dot plot of representative marker genes across cancer subtypes. **(E–F)** UMAP embeddings highlighting strong inter-patient heterogeneity among malignant cells **(G)** Correlation matrix demonstrating transcriptional divergence among malignant subtypes. **(H)** Expression patterns of markers defining proliferative, EMT, mesenchymal-like, ciliated-like, and immunoreactive cancer cell states. See also **Figure S3–S4** and **Data File S5–S6**.

Re-clustering of malignant cells alone revealed strong patient-specific patterns (Fig. 3E–F) that persisted despite computational correction, indicating genuine biological diversity among tumors. Correlation analyses confirmed substantial transcriptomic divergence among malignant cell states (Fig. 3G). Several markers typically associated with the FTE, including *PAX8* and *MUC16*, were broadly expressed across all malignant subtypes (Fig. S4A). Their expression in matched fallopian tube tissues (Fig. S4B–C) further reinforced the FT origin of most tumors. Ciliated-like malignant cells expressed TUBB4B, a hallmark of fallopian tube ciliated epithelium (*22*), consistent with retention of lineage features (Fig. S4A–B). EMT cells exhibited high levels of *MUC16* and *ARGLU1* (Figs. 3H, S4D), whereas mesenchymal-like cells expressed matrix-associated genes such as *COL1A1, COL1A2,* and *VCAN* (Fig. 3H), reflecting tumor core structural phenotypes. Immunoreactive tumor cells expressed *CLU, CD74, HLA-DRA*, and related immune-interaction genes (Figs. 3H, S4C–D; Data File S6). Collectively, these analyses demonstrate that HGSOC exhibits striking inter-patient variability and comprises diverse malignant subpopulations spanning proliferative, ciliated-like, EMT-like, mesenchymal-like, and immunoreactive states. These findings establish a foundation for linking specific malignant phenotypes to the differentiation hierarchy of normal fallopian tube epithelial cells in subsequent analyses.

### The FTE progenitor cells as the most likely origin of HGSOC

Multiple lines of evidence, including our bulk RNA-seq findings, support the concept that HGSOC arises from FTE (*21, 35*). Yet the specific epithelial populations that deviate from normal lineage progression and initiate carcinogenesis have not been firmly established. To address this question, we integrated 11,028 malignant cells from HGSOC samples with 34,267 epithelial cells from normal fallopian tubes, enabling direct comparison of their transcriptional relationships at single-cell resolution (Fig. 4). Using force-directed graph embedding (*36*), we identified 15 distinct cellular clusters encompassing both normal and malignant states (Fig. 4A). Slingshot trajectory analysis suggested that epithelial stem cells (Cluster 11) give rise to three major lineages: mature secretory cells (Cluster 1), ciliated cells (Cluster 3), and malignant cells (Clusters 12 and 13). PAGA connectivity mapping supported these lineage relationships, revealing strong transitions from stem/progenitor clusters toward both normal and cancerous endpoints (Fig. 4B). RNA velocity confirmed that epithelial stem cells and progenitor clusters (0, 4, 8, 11) occupy early pseudotime states and generate trajectories that culminate in either differentiated FTE cell types or HGSOC cell states (Fig. 4C). Global pseudotime analysis demonstrated continuity across all epithelial and cancerous cells, with malignant clusters emerging as late-stage branches diverging from the stem–progenitor axis (Fig. 4D).

**Figure 4.**
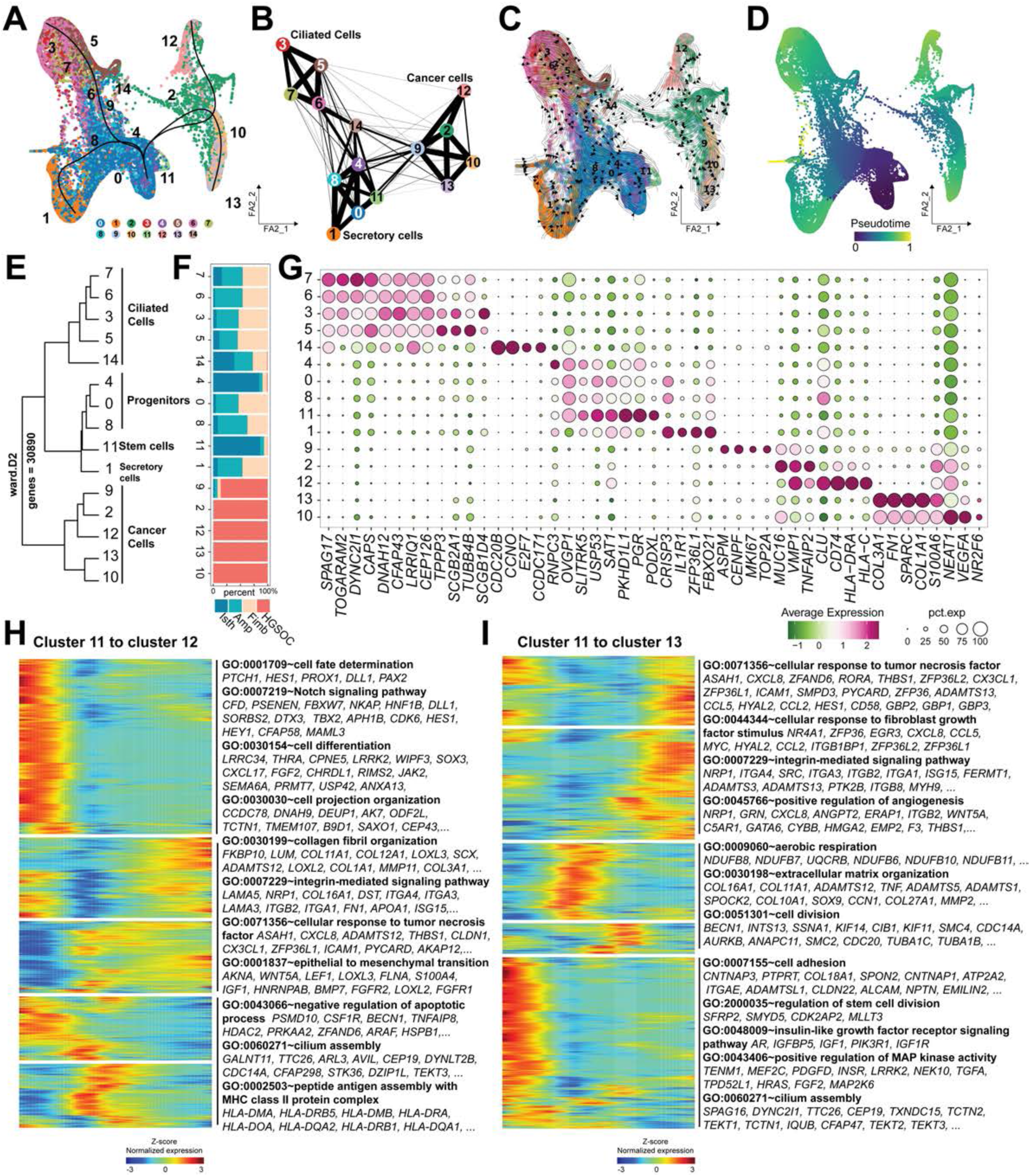
Lineage relationships linking fallopian tube epithelial progenitors to HGSOC malignant states. **(A)** Force-directed graph layout showing 45,295 combined epithelial and cancer nuclei, identifying 15 integrated clusters. **(B)** PAGA analysis reveals three major lineages from epithelial stem cells: secretory cells, ciliated cells, and malignant cells. **(C)** RNA velocity projection demonstrating early progenitors giving rise to either differentiated epithelial states or cancer trajectories. **(D)** Pseudotime analysis showing malignant clusters emerging as late branches from the progenitor axis. **(E)** Hierarchical clustering divides clusters into ciliated, secretory, and cancer cell compartments. **(F–G)** UMAP overlays illustrating that a subset of malignant cells co-cluster with epithelial progenitors, indicating shared transcriptional programs. **(H–I)** Pseudotime heatmaps depicting temporal gene expression patterns along immunoreactive (H) and mesenchymal-like (I) malignant trajectories. See also **Data File S7–S8**.

Hierarchical clustering grouped the 15 clusters into three major compartments: ciliated, secretory, and cancer cell populations (Fig. 4E). Consistent with developmental positioning, progenitor clusters (4, 0, 8) grouped within the secretory compartment alongside epithelial stem cells (Cluster 11) and mature secretory cells (Cluster 1). Although annotated as a ciliated subtype, Cluster 14 exhibited gene expression features and trajectory placement consistent with malignant identity (Figs. 4B–D, 4F–G). Similarly, a subset of malignant cells co-clustered with FTE progenitors (Cluster 4), indicating shared transcriptional programs between early progenitors and cancer-initiated states. Among cancer clusters, we identified four major malignant phenotypes: mesenchymal-like (Clusters 10; *VEGFA*⁺*/NEAT1*⁺), EMT-like (Cluster 13; *SPARC*⁺/*COL1A*⁺), proliferative (Cluster 9; *TOP2A*⁺*/ASPM*⁺), and immunoreactive (Cluster 12; *CD74*⁺*/HLA-DRA*⁺). Notably, proliferative cancer cells incorporated nuclei from fimbriae, ampulla, and isthmus, reflecting strong lineage continuity between progenitors across anatomical regions and the malignant proliferative state (Fig. 4F).

To determine how progenitors diverge toward malignant phenotypes, we analyzed pseudotime progression along two tumor-forming trajectories. From epithelial stem cells to immunoreactive cancer cells (Cluster 12), we observed early downregulation of Notch signaling and cell fate-regulating genes, followed by upregulation of antiapoptotic pathways, cilium assembly programs, and MHC class II antigen processing, consistent with immune-interacting tumor states. Late pseudotime was marked by increased expression of EMT-associated and collagen-organizing genes (Fig. 4H; Data File S7). Along the trajectory leading to mesenchymal-like cancer cells (Cluster 13), early pseudotime reflected loss of stemness-associated transcripts, while late pseudotime showed activation of angiogenesis, integrin signaling, and fibroblast growth pathways, signatures characteristic of proliferating, invasive tumor cells (Fig. 4I; Data File S8). These integrated analyses point to *OVGP1*⁺*/RNPC3*⁺ progenitor cells, particularly those enriched in the fimbrial region, as the epithelial population most primed for malignant transformation. Under normal conditions, progenitors differentiate into either secretory or ciliated epithelium. Under carcinogenic stress or chronic inflammatory stimuli, however, these progenitors can instead divert into malignant pathways, giving rise to immunoreactive or mesenchymal-like tumor states. This dual trajectory mirrors the molecular heterogeneity observed across HGSOC subtypes.

### Transcription factor regulatory networks driving the differentiation from FTE progenitors into normal epithelial or HGSOC cells

To identify the transcriptional regulators that govern normal fallopian tube epithelial (FTE) differentiation and malignant transformation, we performed single-cell regulatory network inference and clustering (SCENIC) on combined datasets of pure FTE cells and HGSOC malignant cells (Fig. 5A). This analysis defined five major regulon-driven cell states corresponding to stem cells, progenitors, secretory cells, ciliated cells, and HGSOC cells. Regulon activity patterns revealed distinct transcription factor (TF) programs defining each cellular state (Fig. 5B). Ciliated cells were characterized by strong activation of *NR2F1, FOXJ1,* and *HOXC4* regulons, canonical drivers of ciliogenesis and epithelial polarization. In contrast, cancer cells exhibited unique activity of regulons such as *TPI1, NR2F6*, and *MAFB*, consistent with metabolic rewiring, proliferative advantage, and tumor-associated immune modulation. Stem cells, progenitors, and mature secretory cells shared substantial overlap in regulon usage, underscoring the fluidity of transitions among these epithelial compartments. Ranking regulons based on specificity (AUC scores) identified *SPDEF* as a prominent regulator in stem and progenitor cells (Fig. 5C). *SPDEF* has previously been implicated in epithelial differentiation and tumor suppression, suggesting a central role in maintaining epithelial identity and restraining aberrant expansion (*37–39*). In contrast, *NR2F6* emerged as one of the most specific and highly active regulators in HGSOC cells (Fig. 5C–D). NR2F6, a member of the nuclear receptor superfamily, is known to influence hormonal response, immune regulation, and oncogenic processes, and has been implicated in multiple types of cancer (*40, 41*). Its high activity across malignant clusters positions it as a potential master regulator of HGSOC cell fate.

**Figure 5.**
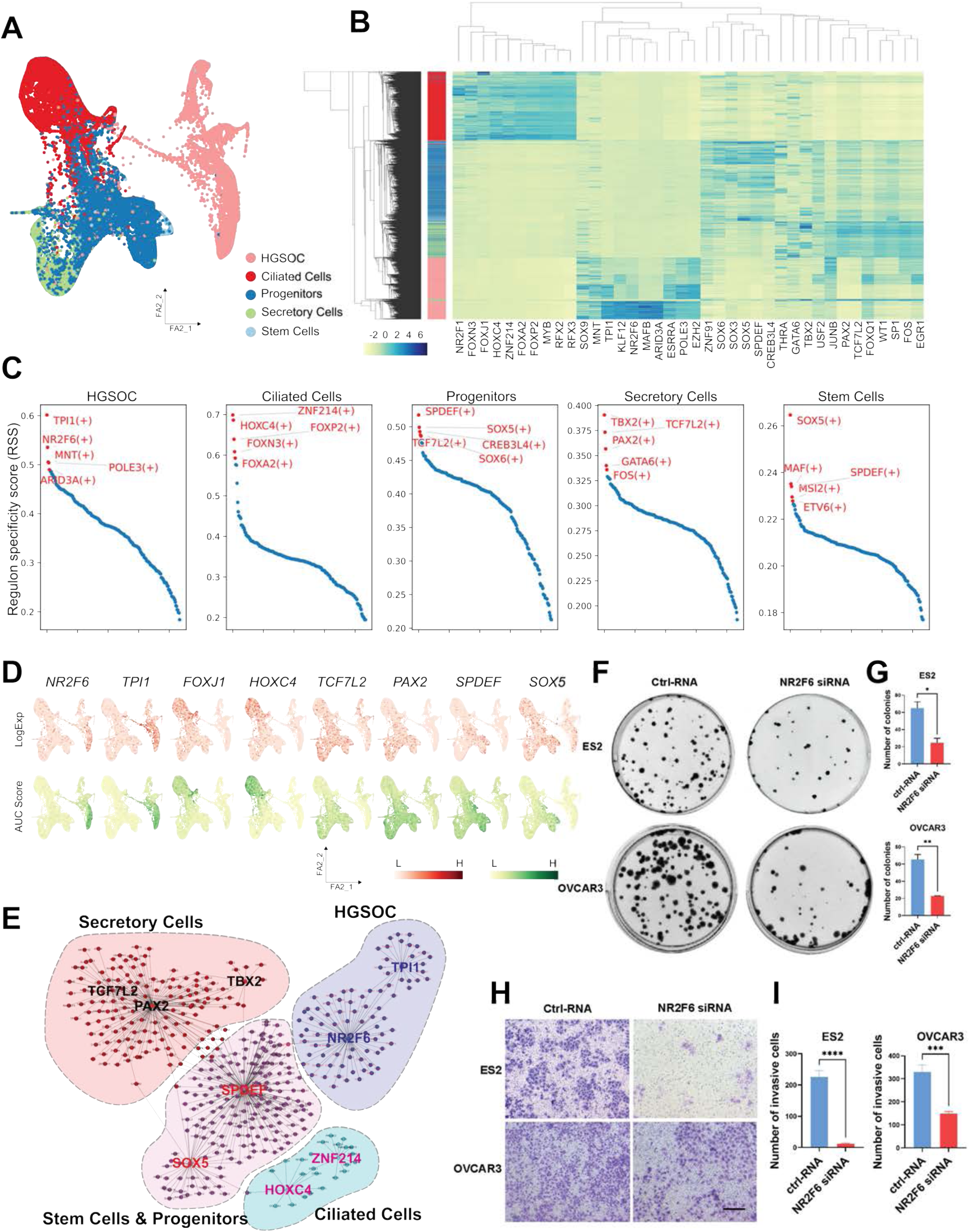
Transcription factor regulatory networks governing normal fallopian tube differentiation and HGSOC malignant transformation. **(A)** Integrative analysis workflow combining normal FTE and HGSOC nuclei for SCENIC regulon inference. **(B)** Heatmap of regulon activity (AUC scores) identifying stem, progenitor, secretory, ciliated, and cancer-specific transcriptional programs. **(C–D)** Ranking of regulons reveals SPDEF as a key stem/progenitor regulator and NR2F6 as a highly specific cancer-associated regulator. **(E)** Gene regulatory network map showing lineage-specific TF modules. **(F–G)** NR2F6 knockdown reduces colony formation in ES2 and OVCAR3 cells. **(H–I)** NR2F6 knockdown decreases invasive capacity of ES2 and OVCAR3 cells in Transwell assays. Data presented as mean ± SEM. Student’s t test. *p* < 0.05; p < 0.01; *p* < 0.001.

Gene regulatory network reconstruction further highlighted SOX5 and SPDEF as master regulators in stem and progenitor states; PAX2, TBX2, and TCF7L2 in mature secretory cells; HOXC4 and ZNF214 in ciliated cells; and NR2F6 and TPI1 in HGSOC cells (Fig. 5E; Fig. S5A). These lineage-specific TF modules delineate the molecular pathways that guide normal epithelial differentiation and, when dysregulated, redirect progenitors toward malignant trajectories. To functionally validate key findings from SCENIC, we knocked down NR2F6 in two human ovarian carcinoma cell lines (ES2 and OVCAR3). NR2F6 suppression significantly reduced colony formation in both lines, indicating reduced proliferative capacity (Fig. 5F–G). Transwell invasion assays revealed a marked decrease in invasive potential following NR2F6 knockdown (Fig. 5H–I), demonstrating that NR2F6 promotes both proliferation and invasion, key hallmarks of HGSOC progression. Together, these data identify distinct transcriptional regulators governing FTE differentiation and HGSOC development, spotlighting NR2F6 as a potential driver of malignant transformation and an attractive therapeutic target.

### DNA copy number variations in HGSOC

Genomic instability is a defining feature of high-grade serous ovarian cancer (HGSOC). To characterize large-scale chromosomal alterations associated with malignant transformation, we inferred DNA copy number variations (CNVs) from snRNA-seq data using inferCNV (*42, 43*), leveraging normal fallopian tube epithelial (FTE) cells as the reference population (Fig. 6A). This approach enabled the detection of extensive chromosomal deletions and amplifications across malignant cells, consistent with the well-documented genomic chaos that typifies HGSOC.

**Figure 6.**
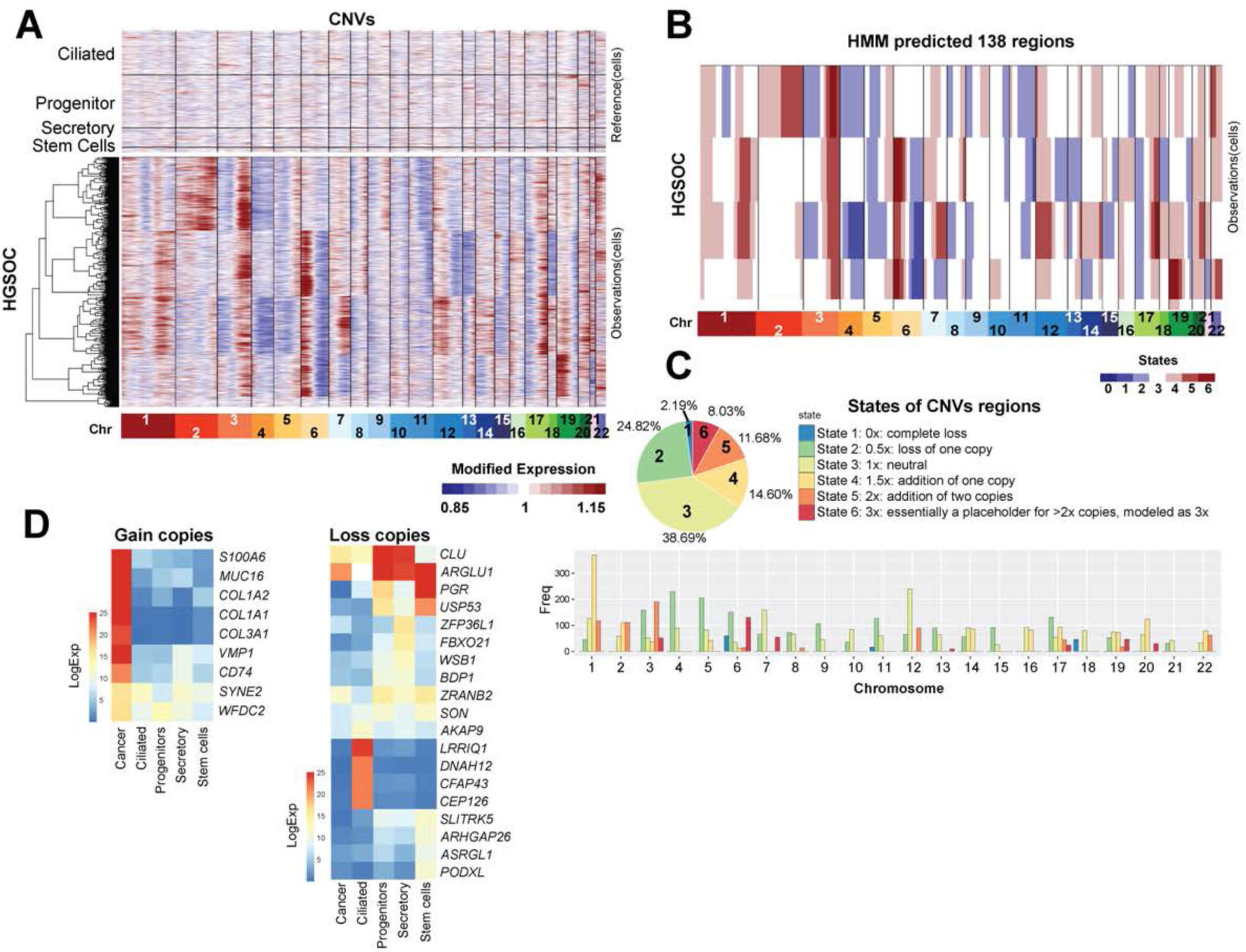
Genome-wide DNA copy number variation (CNV) landscape in HGSOC. **(A)** Overview of inferCNV workflow using FTE epithelial cells as the reference population. **(B)** Hidden Markov Model–based classification of copy number states (0x, 0.5x, 1x, 1.5x, 2x, ≥3x). **(C)** Genome-wide CNV heatmap showing chromosomal losses and gains across malignant cells. **(D)** CNV-driven gene expression biases, highlighting amplified oncogenes (S100A6, MUC16, COL1A1/2, CD74, WFDC2) and genes reduced through copy loss. See also **Data File S9**.

Using a Hidden Markov Model (HMM), we identified 138 CNV regions classified into six states reflecting complete loss (0x), single-copy loss (0.5x), neutral state (1x), single-copy gain (1.5x), two-copy gain (2x), and high-level gain modeled as ≥3 copies (3x) (Fig. 6B; Data File S9). Across all malignant nuclei, ∼39% of genomic regions were copy-neutral, 34% exhibited gains, and 27% displayed losses. Although CNVs were distributed across the genome, Chromosome 1 harbored the highest number of events, while Chromosome 6 displayed the most pronounced amplitude of alteration (Fig. 6C).

Integration of CNV profiles with gene expression revealed that gene dosage effects directly influence many hallmark HGSOC genes. Frequently amplified genes, including *S100A6, MUC16, COL1A1, COL1A2, CD74,* and *WFDC2*, showed corresponding overexpression in malignant clusters (Fig. 6D). Conversely, reduced expression of genes such as *CLU, PODXL,* and *PGR* aligned with copy number loss. These gene-level dosage effects likely contribute to the biased differentiation of progenitor cells toward malignant fates rather than terminally differentiated epithelial states. Collectively, these analyses reveal widespread CNVs in HGSOC and suggest that copy number-driven transcriptional reprogramming reinforces cancer-promoting pathways. These genomic alterations may underlie the divergence of progenitors from normal epithelial maturation trajectories, thereby fostering malignant transformation.

### Pivotal role of collagen signaling in the tumor microenvironment of HGSOC

Extensive interactions between malignant cells and surrounding stromal populations shape the tumor microenvironment (TME) of HGSOC. To delineate these interactions, we examined ligand-receptor networks among cancer cell subtypes, cancer-associated fibroblasts (CAFs), endothelial cells, and macrophages. CAFs emerged as the dominant source of outgoing signals and the most influential regulators of intercellular communication within the TME (Figs. S5B–C). Among all signaling pathways, collagen signaling surfaced as the most prominent extracellular matrix (ECM) regulatory axis. Collagen ligands and their cognate receptors showed extensive cross-talk between CAFs and malignant cells, consistent with collagen’s established roles in matrix remodeling, cellular adhesion, EMT induction, and tumor progression (*44, 45*) (Figs. S5D–E). Multiple collagen-encoding genes, including *COL1A1, COL1A2, COL4A1, COL4A2, COL6A1, COL6A2,* and *COL6A3*, were strongly upregulated in mesenchymal-like and ciliated-like cancer subtypes as well as in CAFs (Fig. S5F), indicating coordinated ECM restructuring. Integrin receptors ITGAV and ITGB8, which form key binding pairs for fibronectin and collagen, were co-expressed in both CAFs and malignant cells. This bilateral expression suggests a feedback loop in which CAFs and tumor cells mutually reinforce an ECM-rich, pro-invasive niche. These interactions likely facilitate tumor cell migration, survival under mechanical stress, and the establishment of metastatic sites. Together, these findings highlight collagen signaling as a central orchestrator of stromal-tumor communication in HGSOC. The prominence of collagen-integrin networks within the TME suggests potential therapeutic opportunities to target CAF-tumor signaling circuits and disrupt ECM remodeling, immunosuppression, and tumor progression.

## DISCUSSION

High-grade serous ovarian carcinoma (HGSOC) remains the most lethal gynecologic malignancy, mainly because its earliest cellular precursors are incompletely understood. Although accumulating evidence implicates the fallopian tube epithelium (FTE), particularly the distal fimbrial region, as the predominant site of origin for many HGSOCs (*46–49*), the specific epithelial populations most vulnerable to malignant transformation have been challenging to define. By integrating bulk RNA sequencing of normal and malignant tissues with high-resolution single-nucleus transcriptomics of cancer-free fallopian tubes and primary HGSOC tumors, this study reconstructs the epithelial hierarchy of the human fallopian tube and identifies epithelial progenitors as the populations most transcriptionally and developmentally aligned with malignant trajectories.

Across all donors, we identify a rare LGR5⁺/PGR⁺ basal cell population that functions as a putative epithelial stem cell pool. These cells give rise to a large, widely distributed population of OVGP1⁺/RNPC3⁺ progenitors, which, in turn, generate both secretory and ciliated epithelial lineages (Fig. 7). This stem-progenitor architecture refines earlier models that primarily emphasized mature secretory cells as the likely cells of origin for HGSOC (*21, 29, 46, 50, 51*). Our integrated analysis demonstrates that it is the progenitor state, not terminal secretory or ciliated states, that shows the strongest transcriptomic and developmental continuity with multiple malignant phenotypes, including immunoreactive, proliferative, and mesenchymal-like tumor states. The transcriptional plasticity, proliferative potential, and intermediate differentiation state of these progenitors may collectively underlie their susceptibility to malignant deviation.

**Figure 7.**
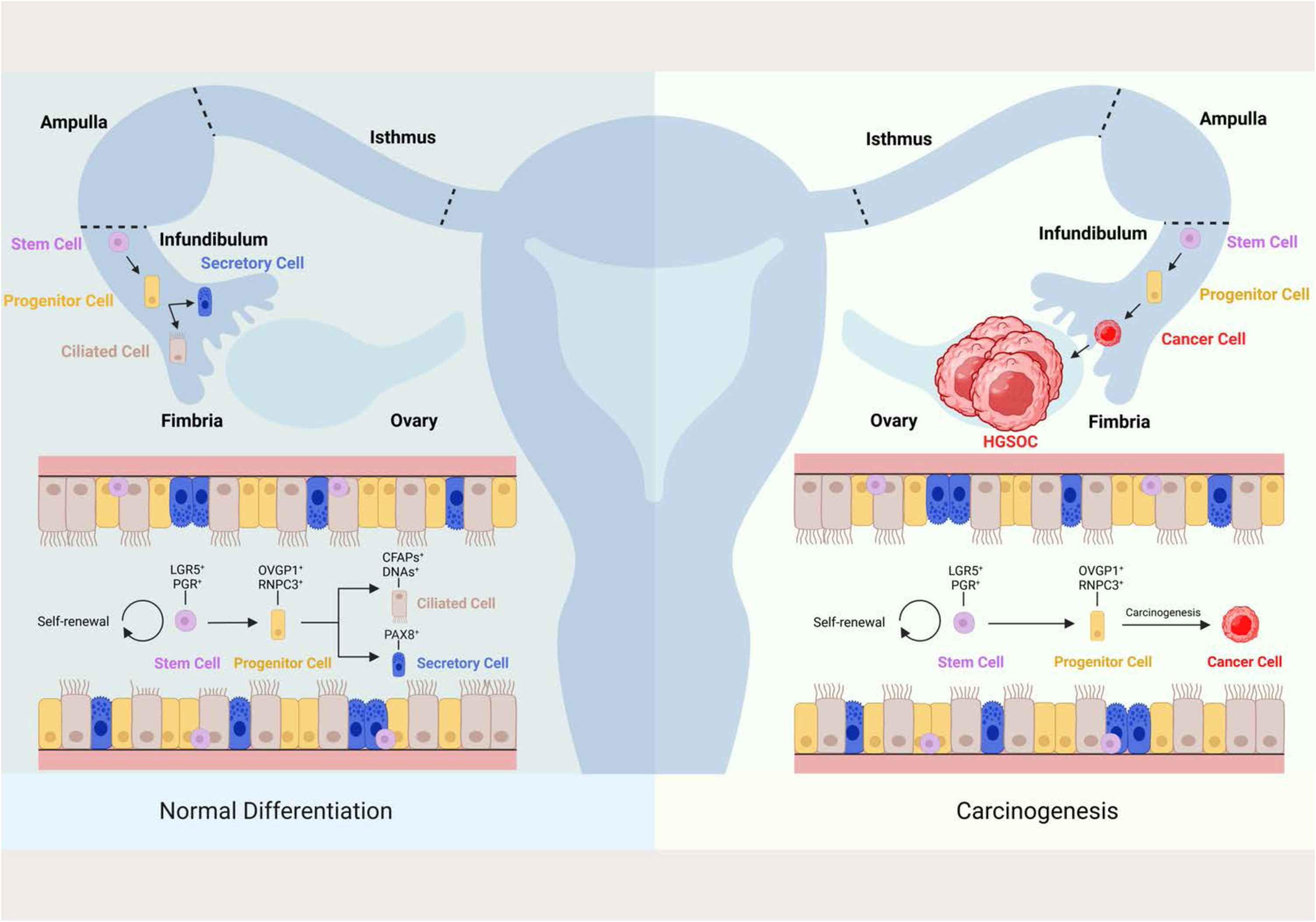
Collagen signaling networks orchestrating stromal–tumor communication in HGSOC. **(A–B)** Ligand–receptor interaction networks showing CAFs as the dominant source of outgoing signals and major regulators of the tumor microenvironment. **(C)** Selected ligand–receptor interaction patterns across malignant and stromal cell types. **(D)** Collagen signaling network indicating extensive cross-talk between CAFs and malignant cells. **(E)** Violin plots of collagen-related gene expression (COL1A1, COL1A2, COL4A1/2, COL6A1/2/3) across major tumor and non-tumor cell subtypes. See also **Figure S5**.

Transcription factor network inference provides additional mechanistic insight. SPDEF emerges as a key regulator of progenitor identity, whereas NR2F6 is highlighted as a candidate driver of malignant lineage divergence. These regulons illustrate how deviations from the normal progenitor program may initiate transcriptional drift toward tumor-like states. Yet transcriptional reprogramming alone does not fully explain the sData File separation between normal epithelial lineages and malignant phenotypes. HGSOC is characterized by pervasive genomic instability, and our inferred DNA copy number variation (CNV) profiles reveal recurrent chromosomal gains and losses that align with oncogenes and malignant regulons. These genomic alterations likely reinforce the lineage deviations identified at the transcriptional level, consolidating malignant identity as tumor trajectories diverge from their normal developmental paths.

The spatial distribution of epithelial cell types along the fallopian tube provides important context for understanding their differential susceptibility to oncogenesis. Our snRNA-seq analyses show that LGR5⁺/PGR⁺ stem cells are most enriched in the isthmus, whereas OVGP1⁺/RNPC3⁺ progenitors form a much larger and more evenly distributed epithelial population across the ampulla and fimbria. Immunofluorescence staining corroborates this pattern, revealing that progenitors occupy the luminal surface throughout distal regions of the tube. These spatial patterns intersect with long-recognized anatomic and physiologic features of the fimbria, including its close proximity to the ovary and its repeated exposure to ovulatory follicular fluid. The fimbria is subjected to cyclical inflammatory microtrauma, bursts of reactive oxygen species, extracellular DNA, and proteolytic enzymes, all of which are known to induce DNA damage and impair epithelial homeostasis (*13, 52–54*). It is, therefore, noData File that the region most frequently harboring serous tubal intraepithelial carcinoma (STIC) lesions also contains a large pool of transcriptionally plastic progenitor cells. The combination of progenitor abundance, ovulatory injury, and a distinct extracellular matrix architecture may create a spatially defined niche in which progenitor states are disproportionately vulnerable to the earliest oncogenic events. Our ligand-receptor analysis further reveals that collagen-integrin signaling between cancer-associated fibroblasts (CAFs) and malignant cells exerts a dominant influence on the HGSOC microenvironment. The fimbrial region’s collagen-rich architecture and dynamic immune milieu may provide additional segment-specific cues that promote malignant divergence once early transcriptional or genomic abnormalities arise in progenitor populations.

Our findings complement and extend existing models of HGSOC initiation. Bulk RNA-seq analyses reveal two transcriptionally distinct tumor groups, one aligning with ovarian surface epithelium and another with fallopian tube epithelium, consistent with a dual-origin model. Earlier models posited that fallopian tube secretory epithelial cells (FTSECs), particularly PAX8⁺/OVGP1⁺ cells, serve as the primary cells of origin for HGSOC (*51, 55, 56*). Still, recent single-cell studies demonstrate substantial secretory heterogeneity and raise questions about which subtypes are most at risk (*57, 58*). Our integrated analysis clarifies this paradigm by showing that progenitors, not mature secretory cells, constitute the most plausible at-risk population (Fig. 7).

Transcriptomic changes along malignant trajectories reveal early tumorigenic features, including anti-apoptotic signaling, metabolic reprogramming, and ECM remodeling, consistent with prior models of serous carcinogenesis (*59–61*). These trajectories converge into well-established HGSOC subtypes with distinct clinical implications (*62, 63*). The tumor microenvironment (TME) plays a central role in shaping malignant progression. CAFs and immune cells regulate ECM dynamics, angiogenesis, and immune suppression. Collagen signaling, driven by CAFs and mesenchymal-like tumor cells, emerged as the dominant ECM pathway in our data, aligning with previous studies demonstrating that collagen deposition, matrix stiffness, and integrin engagement promote tumor invasion, EMT, and metastatic potential (*64, 65*). Interactions between ECM remodeling and tumor-associated macrophages are also well documented (*66–68*), reinforcing the role of the TME in supporting malignant divergence from progenitor states.

Several limitations warrant consideration. First, despite strong developmental continuity between progenitors and malignant states, our data do not constitute direct lineage proof. Definitive demonstration of cellular origin would require in vivo lineage tracing or transformation assays using purified human progenitors, approaches currently not feasible in human tissue. Second, cancer-free fallopian tube samples were obtained from three perimenopausal donors with benign disease; thus, they may not fully capture epithelial diversity across ages or genetic backgrounds. Importantly, tissues from BRCA1/2 mutation carriers, who often harbor early STIC lesions, were unavailable. Future studies should incorporate prophylactic salpingectomy specimens and early lesions. Finally, although our analyses suggest strong microenvironmental influences, spatial transcriptomics and functional assays will be necessary to delineate how stromal and immune contexts shape progenitor vulnerability.

Despite these limitations, our study provides a comprehensive, mechanistically grounded framework linking the normal epithelial hierarchy of the fallopian tube to HGSOC development (Fig. 7). The progenitor-associated markers identified here may inform early detection, risk stratification, and preventive approaches. Targeting lineage-informative regulators, such as NR2F6, or microenvironmental pathways, such as collagen-integrin signaling, may disrupt malignant trajectories rooted in fallopian tube biology. Together, these results support the emerging paradigm that fallopian tube progenitors represent biologically and anatomically plausible epithelial intermediates predisposed to early serous carcinogenesis, offering a foundation for future efforts to intercept HGSOC before clinical manifestation.

## MATERIALS AND METHODS

### Study Design

This study was designed to define the epithelial hierarchy of the human fallopian tube and to determine which epithelial populations show the strongest transcriptional continuity with high-grade serous ovarian carcinoma (HGSOC). We combined bulk RNA sequencing of normal fallopian tubes, ovaries, and primary HGSOC samples with single-nucleus RNA sequencing (snRNA-seq) of cancer-free fallopian tubes and primary HGSOC tissues. Cancer-free fallopian tubes were obtained from perimenopausal donors undergoing gynecologic surgery for benign indications, and HGSOC samples were collected at the time of debulking surgery. No randomization or blinding was applicable to tissue acquisition because the samples derive from human clinical specimens. All snRNA-seq datasets were processed using standardized pipelines, integrated across batches, and analyzed to identify epithelial cell states, developmental trajectories, and inferred transcription factor regulons. Findings from snRNA-seq were validated using bulk RNA-seq comparisons, immunofluorescence staining, pathway analyses, and ligand–receptor modeling. The sample sizes reflect the availability of high-quality human tissues and are consistent with similar single-cell transcriptomic studies. The study was exploratory and mechanistic in nature, aimed at establishing a cellular framework for early serous carcinogenesis rather than testing a prespecified hypothesis.

### Single-Nucleus RNA Sequencing (snRNA-seq)

Single nuclei were isolated from frozen tissues following the **1**0x Genomics Chromium Next GEM v3.1 protocol. Nuclei were encapsulated, barcoded, and amplified to construct 3′ gene expression libraries, which were sequenced to approximately 200 million reads per sample on an Illumina NovaSeq platform (LC-BIO). Data were demultiplexed using the mkfastq function in Cell Ranger v7.1.0, aligned to the hg38 reference provided by 10x Genomics, and quantified using cellranger count (*69*). Secondary data analyses were performed in Python with Scanpy (v1.8.1) (*70*) or in R (v.4.1.0) with the Seurat (v.4.3.0) (*71*), SeuratObject (v4.1.3), SeuratWrappers (v0.3.2) (*71*), tidyverse (v.1.3.2) (*72*), destiny(v3.10.0) (*73*), pheatmap (v1.0.12) packages. Briefly, outputs of cellranger were first loaded into R by using the Read10X function in Seurat, and Seurat objects were built from each sample. Nuclei expressing fewer than 200 genes, more than 7,000 genes, or more than 15% mitochondrial reads were excluded. Data were processed in Seurat using SCTransform for normalization (*74*). Batch effects were corrected using Harmony and LIGER (*34*), and dimensionality reduction was performed using the top 3,000 highly variable genes and the first 20 principal components. Clustering was performed using shared nearest-neighbor modularity optimization, and clusters were annotated based on canonical marker gene expression. Differential gene expression was evaluated using MAST or Seurat’s ROC-based classifier (avgLog_2_FC > 0.25, p < 0.01). Lineage trajectories were reconstructed using PHATE (*75*), diffusion maps (*70, 73*), dpt (*73*), and PAGA (*76*), providing a continuous representation of epithelial and malignant differentiation.

More detailed methods are available in the supplementary material file.

### Statistical Analysis

All statistical analyses were performed using R, Python, Seurat, Scanpy, and associated packages. For snRNA-seq, quality filtering thresholds, normalization methods, dimensionality reduction, integration strategies, and clustering parameters were applied uniformly across samples using edgeR (*77, 78*). Differential gene expression between clusters or conditions was calculated using the Wilcoxon rank-sum test with Benjamini–Hochberg false discovery rate (FDR) correction; genes with adjusted *P* < 0.05 and |log₂FC| ≥ defined thresholds were considered significant. Gene set enrichment analyses used GSEA or Reactome with FDR correction. Inferred copy number variation (CNV) analyses applied a six-state Hidden Markov Model, and RNA velocity analyses used both stochastic and dynamical models for robustness. PAGA connectivity, lineage trajectories, regulon activity (SCENIC), and ligand–receptor interactions (CellChat/NATMI) were evaluated using default or recommended settings. Sensitivity analyses comparing alternative normalization and integration strategies (SCTransform, log-normalization, Harmony, LIGER, Seurat Integration) confirmed the stability of clustering, velocity patterns, and PAGA structure. No statistical methods were used to predetermine sample size due to reliance on human clinical specimens. All statistical tests were two-sided unless otherwise specified. Quantification of immunohistochemistry and immunofluorescence data was conducted using two-tailed Student’s t-tests. Data are presented as mean ± SEM, and p-values <0.05, <0.01, and <0.001 indicate increasing levels of significance.

## List of Supplementary Materials

Materials and Methods

Figs. S1 to S5

Data File s S1 to S2

Data files S1 to S9 (Excel files)

References 1-9 (numbers for references only cited in SM)

## ACKNOWLEDGMENTS

We thank LC-BIO Co., Ltd. (Hangzhou, China) for performing the Chromium 10× Genomics single-nucleus RNA-sequencing and Genedenovo Biotechnology Co., Ltd. (Guangzhou, China) for conducting the Illumina NovaSeq 6000 bulk RNA-seq.

## Funding

Startup funds from The Lundquist Institute for Biomedical Innovation at Harbor-UCLA Medical Center (WY).

Internal research funding from China Medical University (QL).

Startup funds from the Fourth Affiliated Hospital and International School of Medicine, Zhejiang University (KC).

## Competing Interests

Authors declare that they have no competing interests.

## Data and Code Availability

Bulk RNA-seq and single-nucleus RNA-seq (snRNA-seq) datasets have been deposited in the NCBI Gene Expression Omnibus under accession number GSE223426. All custom analysis scripts, computational workflows, and software settings used in this study are openly accessible at https://github.com/chengkeren/fallopiantube, and no restrictions apply to their use. Software versions, computational dependencies, and scripts used in data processing and analysis are documented comprehensively in the GitHub repository (https://github.com/chengkeren/fallopiantube).

## Notes

### Competing Interest Statement

The authors have declared no competing interest.

### Summary of Updates

The main text is now more concise and accurate, and the discussion is more relevant.

