## Supplemental Figures and Tables for "Integrated Human Transcriptomics Identifies Fallopian Tube Progenitors as Plausible Precursors of High-Grade Serous Ovarian Cancer"

**Supplementary Materials include:**

1) Materials and Methods
2) Figs. S1 to S5
3) Tables S1 to S2
4) Data Files S1 to S9

**MATERIALS AND METHODS**

**Human Tissue Collection**

Human fallopian tube and ovarian cancer tissues were collected with informed consent under protocols approved by the Institutional Review Board of the Cancer Hospital of China Medical University and Liaoning Cancer Hospital (Protocol #20220245). For bulk RNA sequencing, high-grade serous ovarian cancer (HGSOC) tissues were obtained from ten patients undergoing cytoreductive surgery. Normal ovaries and fallopian tubes were isolated from five individuals undergoing hysterectomy with bilateral salpingo-oophorectomy for benign uterine conditions. For snRNA-seq, 13 samples from 9 donors were processed, including 5 with benign disease and 4 with confirmed HGSOC. Clinical details for all donors are presented in Tables 1 and 2.

**Donor selection rationale**
Cancer-free fallopian tube samples used for snRNA-seq were obtained from three perimenopausal women undergoing hysterectomy and bilateral salpingo-oophorectomy for benign uterine conditions. Perimenopausal donors were selected intentionally to maximize epithelial cellularity and preserve intact mucosal architecture while avoiding the extensive epithelial atrophy frequently observed in postmenopausal tissues. All donors had macroscopically normal adnexa and no personal or family history suggestive of hereditary cancer syndromes. Although this sample size is limited by the availability of high-quality surgical specimens suitable for nuclei isolation, epithelial subtype composition was highly consistent across all donors.

**Fallopian tube segmentation and sampling**
For single-nucleus RNA sequencing, fallopian tubes were dissected into three anatomical regions, **isthmus**, **ampulla**, and **fimbria,** by an experienced gynecologic pathologist within 1–2 hours of surgical removal. Segments were defined using standard gross anatomical landmarks. The **fimbria** was identified as the fimbriated, fringed distal end of the tube adjacent to the ovary, characterized by a broad infundibulum and delicate, finger-like projections. The **ampulla** was recognized as the widened, tortuous mid-portion containing prominent mucosal folds and a relatively thin muscular wall. The **isthmus** was defined as the narrow, straight, proximal segment near the uterine cornua, with a thicker muscular layer and a reduced luminal diameter. After visual confirmation of orientation via the mesosalpinx attachment, approximately 5–10 mm segments were excised from each region, rinsed in ice-cold PBS, and snap-frozen in liquid nitrogen. Care was taken to avoid contamination from adjacent ovarian cortex or uterine tissue.

**Tissue Processing**

Immediately after surgical removal, fresh tissues were rinsed in ice-cold phosphate-buffered saline (PBS). Portions intended for histological analysis were fixed overnight in 10% neutral buffered formalin, whereas samples designated for RNA extraction or single-nucleus isolation were snap-frozen in liquid nitrogen and stored at –80°C. Fallopian tubes were dissected under a stereomicroscope into three anatomical regions: isthmus, ampulla, and fimbria, which were rinsed in PBS, snap-frozen, and stored at –80°C until further processing.

**Bulk RNA-seq Library Preparation and Sequencing**

Total RNA was extracted using TRIzol (Invitrogen), and RNA integrity was confirmed with an Agilent 2100 Bioanalyzer. Polyadenylated RNA was enriched using oligo(dT) beads, and ribosomal RNA was removed using the Ribo-Zero Magnetic kit (Epicentre). Enriched RNA was fragmented and reverse-transcribed into cDNA. Second-strand synthesis, purification, end repair, poly(A) addition, and Illumina adaptor ligation were performed sequentially. Libraries were size-selected and PCR-enriched prior to sequencing on an Illumina NovaSeq 6000 (PE150) at Genedenovo Biotechnology.

Raw sequencing data were demultiplexed with bcl2fastq2 and adapter-trimmed using Trimmomatic (*1*). Clean reads were aligned to the hg38 human reference genome using STAR (*2*), and gene-level counts were generated using featureCounts (*3*). Weakly expressed genes were filtered using the filterByExpr function in edgeR (*4*). Differential gene expression was assessed using edgeR with thresholds of |log₂ fold change| > 2, p < 0.05, and FDR < 0.05. RPKM values were calculated within edgeR, and downstream visualization, including PCA and hierarchical clustering, was performed in R (v4.1.0).

**RNA Velocity**

RNA velocity analysis was conducted using velocyto to generate spliced and unspliced transcript matrices (*5*). The resulting data were imported into scVelo (v0.2.2) and analyzed using the dynamical model (*6*). Velocity fields were projected onto PHATE and force-directed graph embeddings to infer the directionality and kinetics of transcriptional state transitions.

**SCENIC Regulatory Network Inference**

To infer transcription factor (TF) regulatory networks, we applied the SCENIC pipeline to the integrated FTE and HGSOC datasets. Gene regulatory modules were first inferred using co-expression analyses, followed by motif-based pruning (RcisTarget) to identify direct TF–target relationships (*7*). Regulon activity was quantified using AUC scoring, and cell states were classified based on regulon activity profiles. Gene regulatory networks were visualized in Cytoscape (v3.9.1) (*8*).

**Single-Cell Copy Number Variation Analysis**

Copy number variations (CNVs) were inferred using the inferCNV algorithm (*9*), with fallopian tube stem, progenitor, secretory, and ciliated cells serving as the reference population. Gene expression values were ordered by chromosomal position, smoothed using sliding-window averaging, and truncated at ±3 to limit the influence of outliers. A six-state Hidden Markov Model was used to classify CNV events, corresponding to complete loss (0x), single-copy loss (0.5x), neutral copy number (1x), single-copy gain (1.5x), two-copy gain (2x), and high-level gain (≥3x). CNV landscapes were visualized across all chromosomes.

**Immunohistochemistry (IHC) and Immunofluorescence (IF)**

Formalin-fixed paraffin-embedded tissues were sectioned at 5 µm thickness, deparaffinized, rehydrated, and subjected to citrate-based antigen retrieval. Sections were blocked with 5% normal donkey serum and incubated overnight at 4°C with primary antibodies, including those against LGR5, PGR, Ki67, OVGP1, and CRISP3. Following incubation with secondary antibodies, slides for immunofluorescence were mounted with anti-fade medium, and IHC slides were developed with DAB and counterstained. Images were captured on fluorescence or brightfield microscopes, and quantification was conducted using ImageJ.

**siRNA Transfection**

Two ovarian cancer cell lines, ES2 and OVCAR3, were used in this study. The ES2 ovarian cancer cell line was purchased from Cytion (Cat#305038; Lot# 040324). The OVCAR3 cell line was purchased from ATCC (HTB-161). The quality and reliability of the cell lines have been validated through the company’s in-house and third-party testing, including STR profiling, virus testing, mycoplasma testing, interspecies contamination testing, and bacteria and fungi testing. Ovarian cancer cell lines ES2 and OVCAR3 were seeded in six-well plates at a density of 1×10⁵ cells per well and transfected with NR2F6-specific or negative control siRNAs using jetPRIME®. After six hours, the medium was replaced with complete RPMI 1640 with 10% fetal bovine serum. Knockdown efficiency was assessed by downstream functional assays. siRNA sequences were designed and synthesized by JTSBIO Co., Ltd.

**Colony Formation Assay**

For colony formation, 500 cells were plated per well in six-well plates and cultured for 2–3 weeks, with medium replaced every 2–3 days. Colonies were fixed with 4% paraformaldehyde, stained with crystal violet, imaged using a Bio-Rad imaging system, and counted. The colony formation rate was calculated as the number of colonies divided by the number of seeded cells.

**Transwell Invasion Assay**

Cells were serum-starved for twelve hours, harvested, and resuspended in serum-free RPMI 1640. A total of 5×10⁵ cells were seeded into the upper chamber of Transwell inserts, while the lower chamber contained medium supplemented with 20% serum. After seventy-two hours, cells that had migrated through the membrane were fixed with 4% paraformaldehyde, stained with crystal violet, and imaged for quantification.

**SUPPLEMENTAL FIGURES**

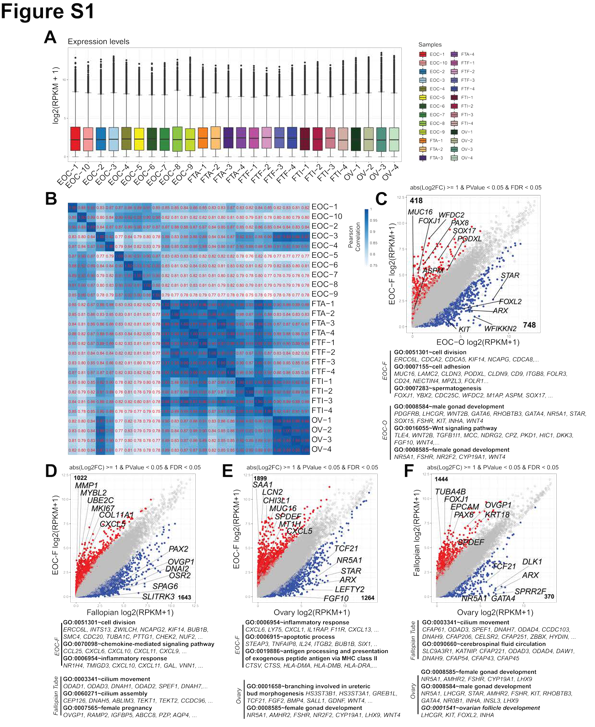

**Figure S1. Transcriptomic correlation among human HGSOCs, fallopian tubes, and ovaries revealed by bulk RNA-seq.**

**(A)** Boxplots showing FPKM-based expression distributions across 26 human samples analyzed by bulk RNA-seq.
**(B)** Heatmap of pairwise Pearson correlation coefficients among all samples.
**(C–F)** Scatterplots of differentially expressed genes (DEGs) comparing:
**(C)** fallopian tube–derived EOC (EOC_F) vs. ovarian-derived EOC (EOC_O);
**(D)** EOC_F vs. fallopian tube;
**(E)** EOC_F vs. ovary;
**(F)** Fallopian tube vs. ovary.
The top enriched Gene Ontology (GO) terms for each comparison are shown.

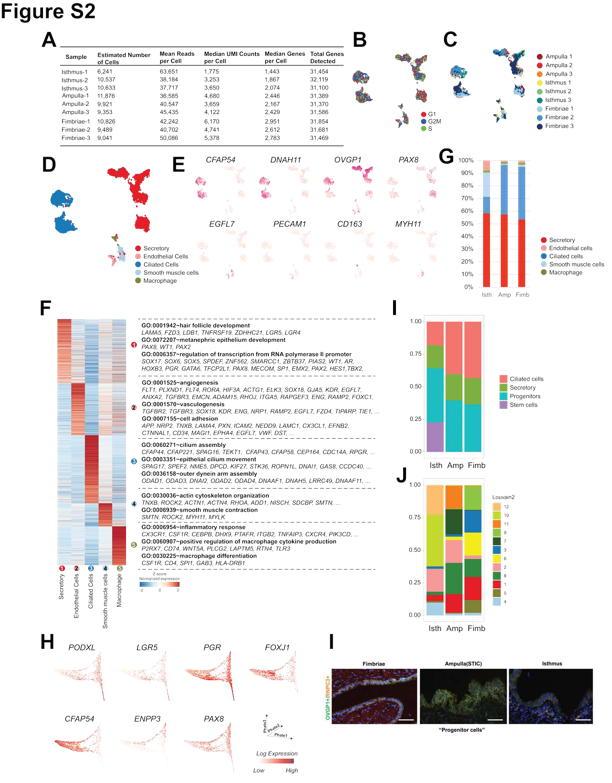

**Figure S2. Major cell subtypes and functional enrichment in human fallopian tubes revealed by snRNA-seq.**

**(A)** Summary of snRNA-seq metrics: 87,920 high-quality nuclei, 36,000–63,000 mean reads per nucleus, and ~2,000 detected genes per nucleus.
**(B–C)** UMAP projections colored by **cell cycle state** (B) and **sample identity** (C) after regression of cell cycle effects.
**(D)** UMAP showing five major fallopian tube cell types: secretory, ciliated, smooth muscle, endothelial cells, and macrophages.
**(E)** UMAP visualizing expression of selected marker genes for major subtypes.
**(F)** Heatmap of top genes and enriched GO terms for each major cell type (Z-score–normalized).
**(G)** Bar graph showing proportional distribution of major cell types across isthmus, ampulla, and fimbria.
**(H)** PHATE trajectory with expression of seven key marker genes.
**(I–J)** Distribution of epithelial subtypes (**I**) and the 12 epithelial subclusters (**J**) across the three fallopian tube segments.
See also **Table S2**.

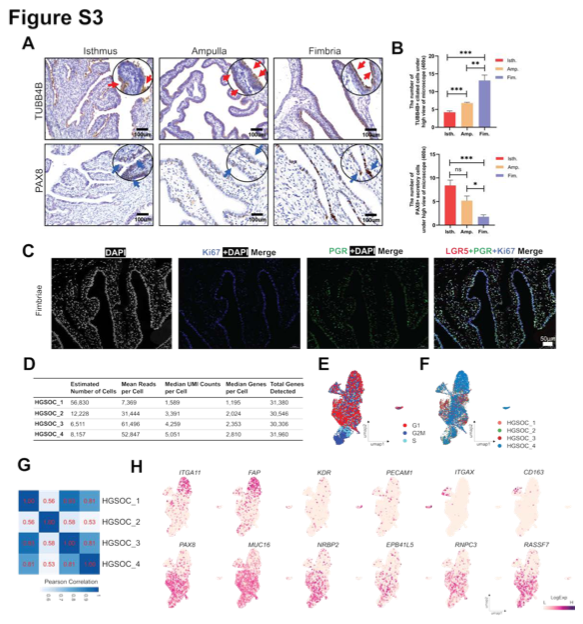

**Figure S3. Prediction and identification of cell subtypes in human fallopian tubes and HGSOC.**

**(A)** Immunohistochemistry (IHC) showing localization of TUBB4B (ciliated cells, red arrows) and PAX8 (secretory cells, yellow arrows) in isthmus, ampulla, and fimbria. Scale bars: 100 µm.
**(B)** Quantification of TUBB4B⁺ and PAX8⁺ cells across segments. Data are mean ± SEM. Student’s t-test. ns ≥ 0.05; *p* < 0.05; **p** < 0.01; ***p*** < 0.001.
**(C)** Immunofluorescence showing co-localization of LGR5, PGR, and Ki67 in fimbrial epithelium. DAPI (white), Ki67 (blue), PGR (green), LGR5 (red). Scale bars: 50 µm.
**(D)** Quality metrics for four HGSOC snRNA-seq samples.
**(E–F)** UMAP projections of HGSOC nuclei colored by **cell cycle state** (E) and **sample identity** (F).
**(G)** Pearson correlation matrix among HGSOC samples.
**(H)** UMAP showing expression of selected HGSOC cell subtype marker genes.

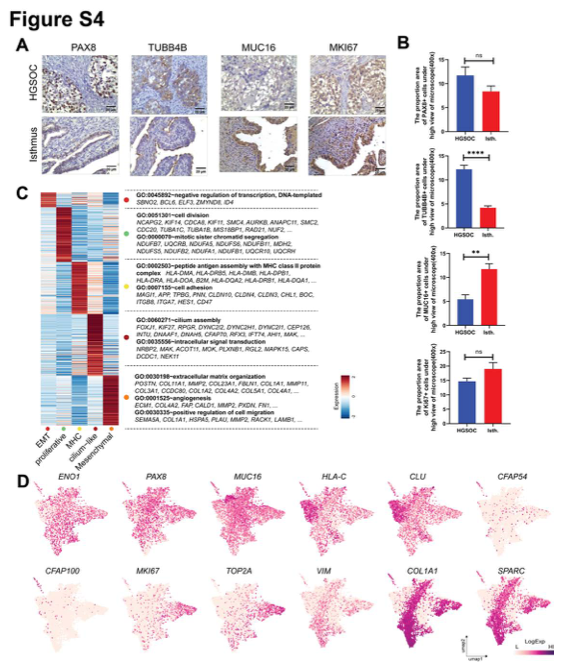

**Figure S4. Specificity and heterogeneity of pure HGSOC tumor cells in snRNA-seq.**

**(A)** IHC showing localization of PAX8, TUBB4B, MUC16, and Ki67 in HGSOC tumors and in isthmus FTE. Scale bars: 20 µm.
**(B)** Quantification of relative abundance of PAX8, TUBB4B, MUC16, and Ki67 in matched tissues. Data are mean ± SEM. Student’s t test; ns ≥ 0.05; **p** < 0.01; **p** < 0.0001.
**(C)** Heatmap showing top genes and enriched GO terms for the five major HGSOC tumor cell subtypes (Z-score-normalized).
**(D)** UMAP projections displaying expression of selected tumor-specific marker genes.
See also **Data File S6**.

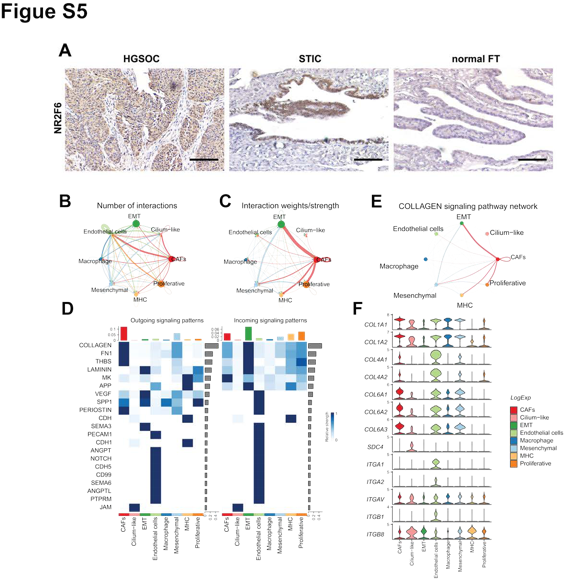

**Figure S5. Intercellular communication in HGSOC.**

**(A–B)** Ligand-receptor interaction networks between cancer and non-cancer subgroups, showing the **number** (A) and **strength** (B) of predicted interactions. Line thickness corresponds to the number of ligand–receptor pairs.
**(C)** Overview of selected outgoing and incoming ligand–receptor signaling patterns across major subtypes.
**(D)** Collagen signaling network showing connectivity between cancer and non-cancer subgroups.
**(E)** Violin plots showing expression of collagen pathway marker genes across tumor and non-tumor subgroups.

**Table S1. Clinical information of the samples used in bulk RNA-seq analyses.**

| Sample (Label) | Age | Reason for surgery^a^ | FIGO staging | Pathology^b^ |
| --- | --- | --- | --- | --- |
| Patient 1 | 51 | EOC | IIIC |  |
| Ovarian cancer tissue (EOC-1) |  |  |  | HGSOC |
| Patient 2 | 61 | EOC | IIB |  |
| Ovarian cancer tissue (EOC-2) |  |  |  | HGSOC |
| Patient 3 | 57 | EOC | IIIC |  |
| Ovarian cancer tissue (EOC-3) |  |  |  | HGSOC |
| Patient 4 | 60 | EOC | IIIC |  |
| Ovarian cancer tissue (EOC-4) |  |  |  | HGSOC |
| Patient 5 | 49 | EOC | IIIC |  |
| Ovarian cancer tissue (EOC-5) |  |  |  | HGSOC |
| Patient 6 | 50 | EOC | IIIC |  |
| Ovarian cancer tissue (EOC-6) |  |  |  | HGSOC |
| Patient 7 | 46 | EOC | IIIC |  |
| Ovarian cancer tissue (EOC-7) |  |  |  | HGSOC |
| Patient 8 | 62 | EOC | IIB |  |
| Ovarian cancer tissue (EOC-8) |  |  |  | HGSOC |
| Patient 9 | 48 | EOC | IVB |  |
| Ovarian cancer tissue (EOC-9) |  |  |  | HGSOC |
| Patient 10 | 62 | EOC | IIB |  |
| Ovarian cancer tissue (EOC-10) |  |  |  | HGSOC |
| Patient 11 | 56 | Endometrial polyps |  |  |
| Ovary (OV-1) |  |  |  | normal |
| Fimbria (FTF-1) |  |  |  | normal |
| Ampulla (FTA-1) |  |  |  | normal |
| Isthmus (FTI-1) |  |  |  | normal |
| Patient 12 | 51 | Hysteromyoma |  |  |
| Ovary (OV-2) |  |  |  | normal |
| Fimbria (FTF-2) |  |  |  | normal |
| Ampulla (FTA-2) |  |  |  | normal |
| Isthmus (FTI-2) |  |  |  | normal |
| Patient 13 | 59 | Hysteromyoma |  |  |
| Ovary (OV-3) |  |  |  | normal |
| Fimbria (FTF-3) |  |  |  | normal |
| Ampulla (FTA-3) |  |  |  | normal |
| Isthmus (FTI-3) |  |  |  | normal |
| Patient 14 | 53 | Hysteromyoma |  |  |
| Ovary (OV-4) |  |  |  | normal |
| Fimbria (FTF-4) |  |  |  | normal |
| Ampulla (FTA-4) |  |  |  | normal |
| Patient 15 | 49 | Hysteromyoma |  |  |
| Isthmus (FTI-4) |  |  |  | normal |

^a^EOC, epithelial ovarian cancer. ^b^HGSOC, high-grade serous ovarian cancer.

**Table S2. Clinical information of the samples used in single-nucleus RNA-seq analyses.**

| Sample | Age | Reason for surgery | Pathology | #Cell | Mean reads/Cell | Median genes/Cell |
| --- | --- | --- | --- | --- | --- | --- |
| Patient 1 | 47 | Hysteromyoma and adenomyosis |  |  |  |  |
| Fimbria |  |  | normal | 10,928 | 41,847 | 2,887 |
| Ampulla |  |  | normal | 11,580 | 37,520 | 2,437 |
| Isthmus |  |  | normal | 6,406 | 62,011 | 1,392 |
| Patient 2 | 46 | Hysteromyoma and adenomyosis |  |  |  |  |
| Fimbria |  |  | normal | 9,517 | 40,582 | 2,560 |
| Ampulla |  |  | normal | 9,924 | 40,534 | 2,128 |
| Patient 3 | 42 | Hysteromyoma |  |  |  |  |
| Isthmus |  |  | normal | 10,531 | 38,206 | 1,837 |
| Patient 4 | 51 | Hysteromyoma and adenomyosis |  |  |  |  |
| Ampulla |  |  | normal | 9,588 | 44,321 | 2,355 |
| Patient 5 | 56 | Hysteromyoma |  |  |  |  |
| Fimbria |  |  | normal | 9,147 | 49,505 | 2,716 |
| Isthmus |  |  | normal | 10,377 | 38,647 | 2,071 |
| Patient 6 |  |  |  |  |  |  |
| OC tissue | 57 | EOC | HGSOC | 11,389 | 36,769 | 1,843 |
| Patient 7 |  |  |  |  |  |  |
| OC tissue | 59 | EOC | HGSOC | 11,334 | 33,924 | 2,070 |
| Patient 8 |  |  |  |  |  |  |
| OC tissue | 50 |  | HGSOC | 6,825 | 58,667 | 2,255 |
| Patient 9 |  |  |  |  |  |  |
| OC tissue | 53 | EOC | HGSOC | 8,157 | 52,847 | 2,810 |

OC, ovarian cancer; EOC, epithelial ovarian cancer; HGSOC: high-grade serous ovarian cancer

**Data Files S1 to S9, see separate Excel Files**

**Data File S1.** Top 1000 variable genes among different samples, related to Figure 1DE.

**Data File S2.** Markers identified across different cell types in human fallopian tube snRNA-seq, as shown in Figure S2F.

**Data File S2.** Top 2000 variable genes along the secretory trajectory, related to Figure 2H.

**Data File S4.** Top 2000 variable genes along the ciliated trajectory, related to Figure 2I.

**Data File S5.** Markers identified in HGSOC snRNA-seq, related to Figure 3D.

**Data File S6.** Markers identified among subtypes of HGSOC cancer, as shown in Figure S4C.

**Data File S7.** Top 200 variable genes along the trajectory from cluster 11 to cluster 12, related to Figure 4H.

**Data File S8.** Top 200 variable genes along the trajectory from cluster 11 to cluster 13, related to Figure 4I.

**Data File S9.** HMM predicted CNV regions, related to Figure 6B.
